# High-throughput identification of MHC class I binding peptides using an ultradense peptide array

**DOI:** 10.1101/715342

**Authors:** Amelia K. Haj, Meghan E. Breitbach, David A. Baker, Mariel S. Mohns, Gage K. Moreno, Nancy A. Wilson, Victor Lyamichev, Jigar Patel, Kim L. Weisgrau, Dawn M. Dudley, David H. O’Connor

## Abstract

Rational vaccine development and evaluation requires identifying and measuring the magnitude of epitope-specific CD8 T cell responses. However, conventional CD8 T cell epitope discovery methods are labor-intensive and do not scale well. Here, we accelerate this process by using an ultradense peptide array as a high-throughput tool for screening peptides to identify putative novel epitopes. In a single experiment, we directly assess the binding of four common Indian rhesus macaque MHC class I molecules – Mamu-A1*001, -A1*002, -B*008, and -B*017 – to approximately 61,000 8-mer, 9-mer, and 10-mer peptides derived from the full proteomes of 82 simian immunodeficiency virus (SIV) and simian-human immunodeficiency virus (SHIV) isolates. Many epitope-specific CD8 T cell responses restricted by these four MHC molecules have already been identified in SIVmac239, providing an ideal dataset for validating the array; up to 64% of these known epitopes are found in the top 192 SIVmac239 peptides with the most intense MHC binding signals in our experiment. To assess whether the peptide array identified putative novel CD8 T cell epitopes, we validated the method by IFN-γ ELISPOT assay and found three novel peptides that induced CD8 T cell responses in at least two Mamu-A1*001-positive animals; two of these were validated by *ex vivo* tetramer staining. This high-throughput identification of peptides that bind class I MHC will enable more efficient CD8 T cell response profiling for vaccine development, particularly for pathogens with complex proteomes where few epitope-specific responses have been defined.

## Introduction

CD8 T cell receptors typically recognize epitopes – linear peptides of 8-11 amino acids – presented by major histocompatibility complex (MHC) class I molecules. These peptides are usually derived from self proteins that under normal circumstances do not induce an immune response. Cancerous cells, transplanted cells, and infected cells, however, present non-self peptides that are recognized as foreign by CD8 T cells (1). CD8 T cell recognition of foreign epitopes, including those derived from viral proteins, along with concurrent recognition of costimulatory molecules on the antigen-presenting cell, leads to T cell activation and production of cytotoxic compounds that trigger apoptosis in the affected cell (2). Polymorphisms in the peptide binding cleft of the MHC glycoprotein determine the variety of peptides each molecule is able to present, and specificity is determined in part by anchor residues in the peptide binding to specific pockets in the binding cleft (3). Identifying which peptides are presented by particular MHC molecules and are likely to elicit epitope-specific CD8 T cell responses is a critical task in designing prophylactic and therapeutic vaccines against infectious diseases and cancers (4, 5).

Characterizing MHC-peptide binding can be a time- and resource-intensive task. Competitive binding assays, which calculate the concentration (IC_50_) at which a peptide of interest outcompetes 50% of a radiolabeled probe peptide for binding to MHC class I molecules, are used to estimate MHC-peptide binding affinity (6). These methods are low-throughput and difficult to scale. Computational tools have been developed that have the potential to improve throughput, enabling prediction of the likelihood a given peptide will bind to a specific MHC molecule. However, these methods exploit our understanding of anchor residues and rely on previously generated peptide binding datasets for pattern recognition, and so are limited to MHC molecules whose binding repertoires have already been well studied, creating a self-fulfilling prophecy of sorts (6, 7). A different computational technique, molecular docking, uses the three-dimensional crystal structure of MHC molecules to predict peptides likely to bind (8, 9). While this method has the benefit of working for both MHC class I and II molecules, it still cannot serve as a substitute for *in vitro* binding assays.

CD8 T cell epitope identification is a similarly involved task, often requiring large quantities of cells exposed *in vivo* to the pathogen of interest. In our previous work identifying CD8 T cell epitopes, we performed IFN-γ ELISPOTs using pools of 15-mer peptides to stimulate peripheral blood mononuclear cells (PBMCs), followed by successive rounds of deconvolution of positive wells to determine stimulatory peptides and minimal optimal epitopes within positive 15-mers (10). This is followed by generation of MHC-peptide tetramers, which stably bind T cell receptors and can be used for cell staining and sorting (11). While successful, these methods are not ideal – using 15-mer peptides, which are too large to bind class I MHC *in vivo*, can lead to false negatives in ELISPOT assays (12). Currently, there is no high-throughput *in vitro* tool for assessing whether shorter peptides are likely to bind to MHC and therefore are more likely to be CD8 T cell epitopes.

Rhesus macaques infected with simian immunodeficiency virus (SIV) and simian-human immunodeficiency virus (SHIV) have long been used as models for HIV infection studies. Rhesus macaque class I MHC class I A (Mamu-A) and class I B (Mamu-B) genes bind and present peptides to CD8 T cells like their HLA-A and HLA-B counterparts in humans. Previous studies have characterized the peptide binding specificity of common Mamu-A and Mamu-B molecules to SIV-derived epitopes, and have identified epitopes that elicit CD8 T cell responses (13–16). Knowledge of these epitopes enables tracking of SIV-specific responses after infection and vaccination (17), and identification of the peptide binding specificities of MHC alleles commonly found in elite controllers has guided the development of epitope-based vaccines against SIV (18). Yet, even in this well-characterized system, T cell epitopes have only been defined for a small minority of the hundreds of known MHC class I alleles, while nearly no T cell epitopes have been defined for other macaque viruses or bacterial pathogens. While considerably more effort has been dedicated to defining T cell epitopes in humans, these efforts are similarly concentrated on a small number of pathogens (e.g., HIV, influenza). Methods that facilitate the high-throughput identification of T cell epitopes, particularly for pathogens with large proteomes where T cells play an essential role in protective immunity (e.g., *Mycobacterium tuberculosis*), are urgently needed.

Ultradense peptide arrays have previously been used as a high-throughput tool for assessing antibody responses to pathogens; here, we repurpose them to identify peptides that bind specific MHC class I molecules. Briefly, up to six million linear peptides are synthesized on a single array chip, molecules of interest – antibodies in serum, or other molecules such as MHC proteins – are allowed to bind to the peptides, and binding affinity is reported in fluorescence intensity units based on the detection of fluorophore-conjugated secondary antibodies bound to the molecules. Our group and others have designed arrays to simultaneously assess IgG and IgM binding to linear pathogen-derived peptides (19, 20). Here, we generated a peptide array with 61,066 unique 8-mer, 9-mer, and 10-mer peptides derived from 82 SIV and SHIV isolates, and measured binding to four rhesus macaque MHC class I molecules, Mamu-A1*001, -A1*002, -B*008, and -B*017, all of which have peptide binding repertoires that have been extensively studied by conventional methods. While we focus our attention here on a particular well-studied strain, SIVmac239, as a way of validating the array, the large capacity of the array gave us the opportunity to generate the same data for several dozen isolates, creating a sizable dataset that allows for comparisons among strains. We tested the top 86 SIVmac239-derived peptides identified by the array as having the highest binding affinity for Mamu-A1*001 for their ability to induce IFN-γ production in peripheral blood mononuclear cells isolated from SIVmac239-infected, Mamu-A1*001-positive rhesus macaques. We demonstrate that peptides that exhibit strong MHC binding on our array frequently correspond to CD8 T cell epitopes that have been identified by conventional assays, and that our high-binding peptides are enriched for putative novel epitopes that elicit CD8 T cell responses. This suggests that the MHC-peptide array can be used to predict immunologically relevant MHC-peptide interactions, offering a high-throughput alternative to traditional methods.

## Materials and Methods

### SIV/SHIV peptidome array design

Peptide sequences were generated from SIV and SHIV amino acid sequences (Supplemental table I). We downloaded 82 SIV and SHIV GenBank files from NCBI and extracted amino acid sequences for all open reading frames into FASTA files. The GenBank SIVmac239 nef protein sequence is truncated and therefore the correct full-length sequence was translated from the full GenBank sequence and added manually. The FASTA files were sent to Roche NimbleGen (Madison, WI; now Nimble Therapeutics) to generate the array. Non-redundant 8-mer, 9-mer, and 10-mer peptide sequences were generated with 1-amino acid offsets, and 5 replicates for each peptide were included on the array in a 12-plex array configuration.

### Amino acid substitution array design

Substitution libraries were designed by introducing all possible single amino-acid substitutions and single-amino-acid deletions for the Gag CM9 epitope (CTPYDINQM) and the Tat SL8 epitope (STPESANL) using all 20 L-amino acids. The library was synthesized in five replicates. For this array, 1785 peptides were synthesized (9 × 21 × 5 + 8 × 21 × 5 = 1785, where 9 and 8 are the peptide lengths, 21 is the number of possible amino acids/deletion at each position, and 5 is the number of replicates).

### Production of MHC molecules

Inclusion bodies were prepared essentially according to the method of Hutchinson, et al (21). In brief, bacterial expression plasmids containing the sequence for human beta-2 microglobulin (β2M) or rhesus MHC class I Mamu-A1*001, Mamu-A1*002, Mamu-B*008, or Mamu-B*017 were transformed into competent BL21DE3 *E. coli* bacteria. Single colonies were grown in 1 L TYP media (16 g Yeast extract, 16 g Tryptone, 5 g NaCl, 1 g K_2_HPO_4_) with appropriate antibiotic markers, induced with IPTG during logarithmic growth phase, and allowed to express protein overnight. Bacterial pellets were incubated with lysis buffer (10 mM Tris pH 8.1, 150 mM NaCl, 10 mM EDTA with DNAse and MgCl_2_). Following sonication, the inclusion bodies were purified in 50 mM Tris pH 8.1 10 mM EDTA 100 mM NaCl buffer with 0.5% Triton X-100, washed in resuspension buffer without Triton-X and the pellet dissolved in denaturing buffer. β2M inclusion bodies were dissolved in 8M Urea with 50 mM Tris pH 8.1, 100 mM NaCl, 10 mM EDTA and 10 mM DTT. MHC heavy chains were dissolved in 6 M guanidine, 50 mM Tris pH 8.1, 100 mM NaCl, 10 mM EDTA with 10 mM DTT. After a Bradford assay to determine the concentration of protein, inclusions were snap frozen in LN2 and frozen at −80°C in aliquots of 30 mg for MHC heavy chain and 10 mg for β2M.

### MHC complex preparation

To prepare MHC I complexes for peptide array binding, 45 μl di-sodium tetraborate/sodium hydroxide pH 10 buffer (Sigma-Aldrich), 10 μl 2 mg/ml β2M, and 60 μg of MHC class I heavy chain stock were mixed in 400 μl water. This mixture was concentrated with AmiconUltra 0.5 ml, 10K centrifugal filter (Millipore) at 12,000 g for 5 min without pre-incubation. Concentrated material was mixed with 400 μl 0.05X pH10 buffer to further dilute urea and 2-ME and concentrated in the same Amicon filter at 12,000 g for 7 min. The concentrated sample was transferred to a 1.5 ml Eppendorf tube, centrifuged at 20,000 g for 5 min to remove precipitated material, and stored at 4°C before binding to the peptide array.

### MHC-peptide binding assay

Peptide synthesis was accomplished through light-directed array synthesis in a Maskless Array Synthesizer (MAS) using an amino-functionalized substrate as previously reported (22). Each peptide replicate were randomly positioned on the array. After synthesis and de-protection, the array was incubated in 0.67 μg/ml Cy5-labeled streptavidin (SA-Cy5) (Amersham), 1% alkali-soluble casein (Novagen) at room temperature for 1 hour to stain technical SA-binding features used for array QC and gridding. After a quick rinse in water, the array was dried by spinning in a microcentrifuge equipped with an array holder and a 12-plex incubation chamber was applied to the array surface. Before loading onto the array, MHC I complexes were 1:6 diluted in 20 mM tris-HCl, pH7.4, 1% BSA and 6 μl aliquots were loaded into 5 chambers of a 12-plex chamber well. An additional chamber was loaded without the MHC I complexes as a control. The loading ports were sealed with port seals (Grace Bio-Labs, Bend, OR) and the array was incubated at 4°C overnight. After incubation the chamber was removed, and slides were rinsed in water and stained by dipping with Alexa Fluor 647 MEM-123 1:1,000 (Novus Biologicals, Centennial, CO). After rinsing in water and drying as described above, the array was scanned to collect fluorescence data. Cy5 fluorescence intensity of the array was measured with an MS200 scanner at resolution 2 μm, wavelength 635 nm. Cy5 fluorescence intensities were extracted using Image Extraction Software.

### Peptide array data analysis

The array data contains the fluorescence intensity, reported in arbitrary signal intensity units, for each peptide sequence replicate for each MHC molecule. We took the base 2 logarithm of the fluorescence intensity and used the median value for each of the unique peptide sequences. Using the median reduces noise induced by the peptide position on the array and reduces the overall variance. We assessed the top 24, 48, 96, and 192 peptides with the highest binding intensities in each virus-MHC class I molecule combination for the presence of known epitopes.

### IFN-γ ELISPOT assay

ELISPOTs were performed in duplicate using frozen rhesus macaque PBMCs isolated from EDTA-treated whole blood. Frozen cells were thawed slowly at room temperature and washed twice in R10 media containing RPMI, 10% FBS, 1% antibiotic/antimycotic, and 1% L-glutamine (Hyclone, Logan, UT). Approximately 100,000 cells in 100 μl of R10 media were layered on precoated monkey IFN-γ ELISpot-Plus plates (Mabtech Inc., Mariemont, Ohio, USA) with a single peptide per well at a final concentration of 1 μM. Peptides were synthesized by GenScript (Piscataway, NJ, USA). Concanavalin A (ConA, 10 μM) was used as a positive control and cells with no peptide were used as a negative control.

### ELISPOT analysis

Developed ELISPOT plates were read with an AID Elispot reader using the IFN-γ setting and a threshold of 100 spots per million cells. ConA wells were counted using custom parameters defined for ConA (Min intensity: 10, Min size: 20, Min gradient: 1, Max intensity: 57, Max size: 295, Max gradient: 50, Basic algorithm settings emphasis: tiny, Algorithm: C); the rest were read using custom IFN-γ parameters (Min intensity: 20, Min size: 40, Min gradient: 1, Max intensity: 254, Max size: 5000, Max gradient: 90, Basic algorithm settings emphasis: small, Algorithm: C). Positive ELISPOT responses were determined as previously described (23).

### Tetramers and tetramer staining

Tetramers conjugated to APC or PE were generously produced by the NIH Tetramer Core Facility using peptides generated by GenScript. Thawed PBMC resuspended to 1 × 10^6^ cells/100 μl of R10 medium were stained with tetramers at a concentration of 5 μg/ml in the dark for one hour at 37°C, and were subsequently stained with CD3 Alexa Fluor 700 (BD Biosciences, San Jose, CA) and CD8 BV421 (BD Biosciences) for 15 minutes at room temperature, washed, and fixed with 2% paraformaldehyde. Samples were run on a BD LSRII flow cytometer within 24 hours and were analyzed with FlowJo software, version 10.5.3.

## Results

### Identification of SIVmac239 peptides that bind MHC class I molecules

We tested binding of purified rhesus MHC class I Mamu-A1*001, -A1*002, -B*008, and -B*017 molecules to the full proteomes of 82 SIV and SHIV isolates (Supplemental table I). Previous studies have identified SIVmac239 epitopes that bind each of the four MHC class I molecules tested here, which enables us to validate the peptide array as a method for screening for putative novel CD8 T cell epitopes (14–16, 24). To determine whether the array identified previously-established SIVmac239 epitopes, we determined the percentage of known CD8 T cell epitopes that were identified in the top 24, 48, 96, and 192 (convenient fractions/multiples of a 96-well ELISPOT plate) highest-binding peptides for each MHC molecule. The top 192 peptides for each of the four MHC molecules encompassed 27-64% of the known T cell epitopes restricted by each molecule (Table I). Figure 1 shows the complete data for SIVmac239 peptides binding to Mamu-A1*001, -A1*002, -B*008, and -B*017, with previously-identified epitopes present in the top 192 peptides (out of 9912 SIVmac239 peptides, or 1.9%) for each molecule indicated.

**Table I:**
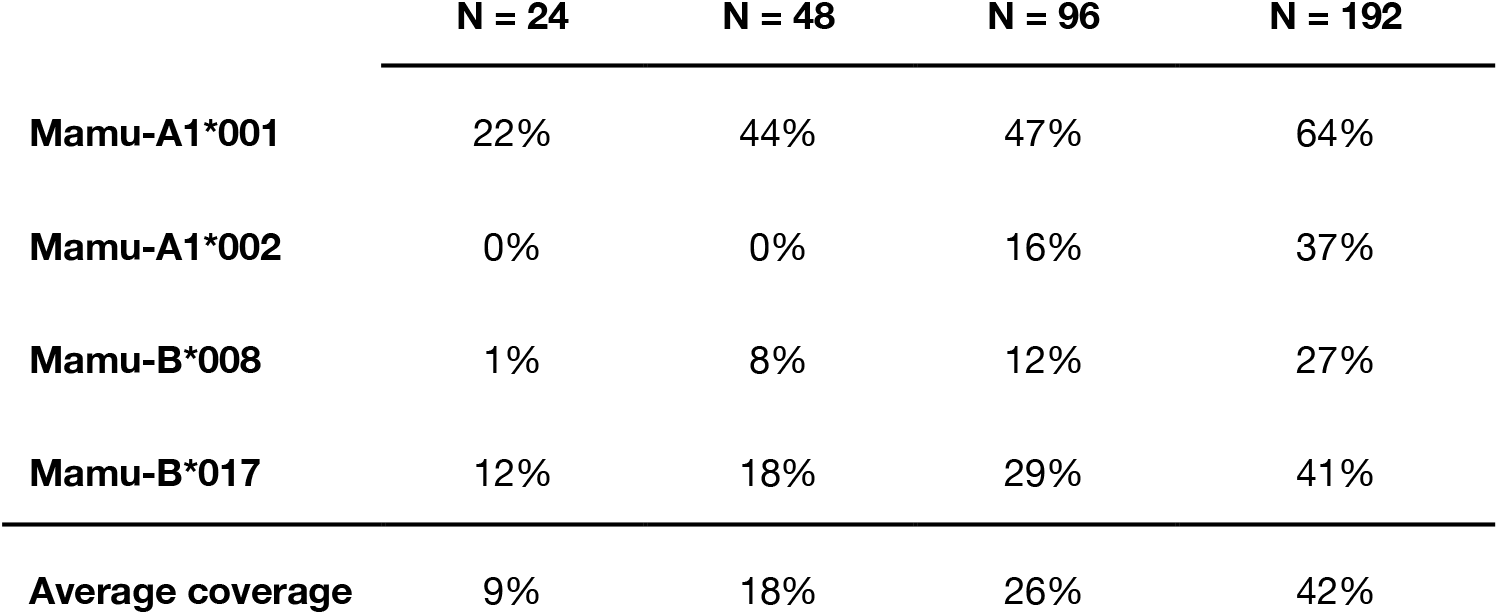
The percentage of known epitopes for each MHC molecule that were found in the top N peptides with high signal intensity.

**Figure 1:**
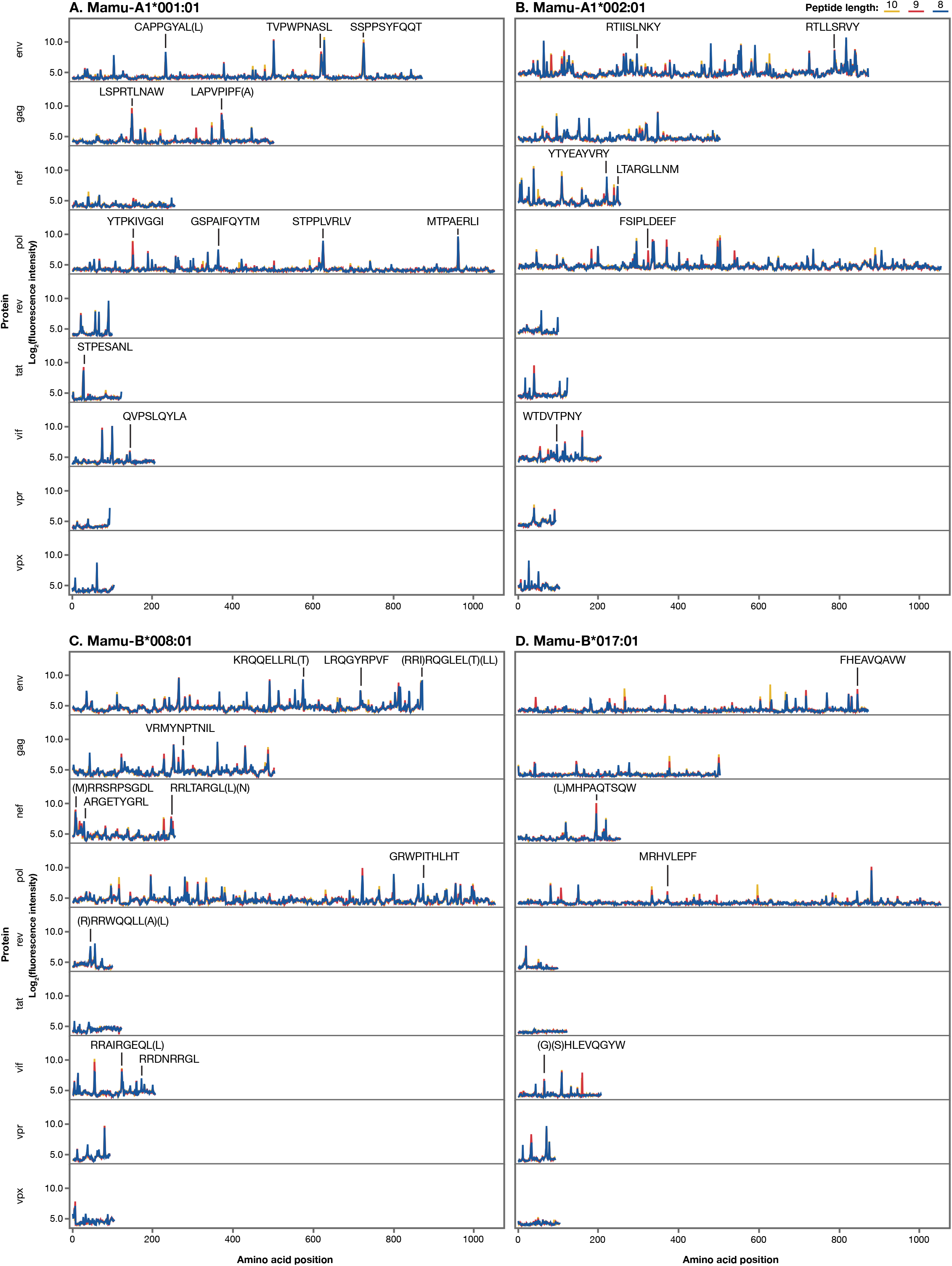
MHC-peptide binding plots. Previously-identified SIVmac239 8-mer, 9-mer, and 10-mer epitopes found in the top 192 peptides for each MHC class I molecule. Parentheticals indicate overlapping epitopes.

### Critical residues for Mamu-A1*001 peptide binding identified through substitution analysis

Peptide anchor residues define the binding motif for a given MHC molecule and enable prediction of other peptides likely to bind. The binding motif for Mamu-A1*001 was previously defined by sequence analysis of peptides bound to the molecule in vitro and by performing competitive binding assays using peptides with amino acid substitutions (24, 25). These studies identified positions two and three (P2 and P3), and the C-terminus, as important for binding. We performed a full substitution analysis at each amino acid for two known immunodominant CD8 T cell epitopes that bind to Mamu-A1*001, Gag CM9 (Gag 181-189) and Tat SL8 (Tat 28-35). We tested the influence of all 20 amino acids, plus an amino acid deletion, at each position on the binding of the peptides to Mamu-A1*001 molecules. As would be expected based on prior studies, substitutions at P2 (threonine) and P3 (proline) for both peptides were generally poorly tolerated and resulted in dramatically decreased signal intensity, indicating reduced binding affinity (Figure 2). At P2, a substitution with serine was tolerated; P3 did not tolerate any substitutions. These results corroborated those found by competitive binding assays using Gag CM9 peptides with single amino acid substitutions (24). A previous study found that L, I, V, M, F, W, Y, and T are tolerated at the C-terminus; however, this study did not test every possible amino acid at that position, and our results suggest that R, K, and H may also be tolerated (24).

**Figure 2:**
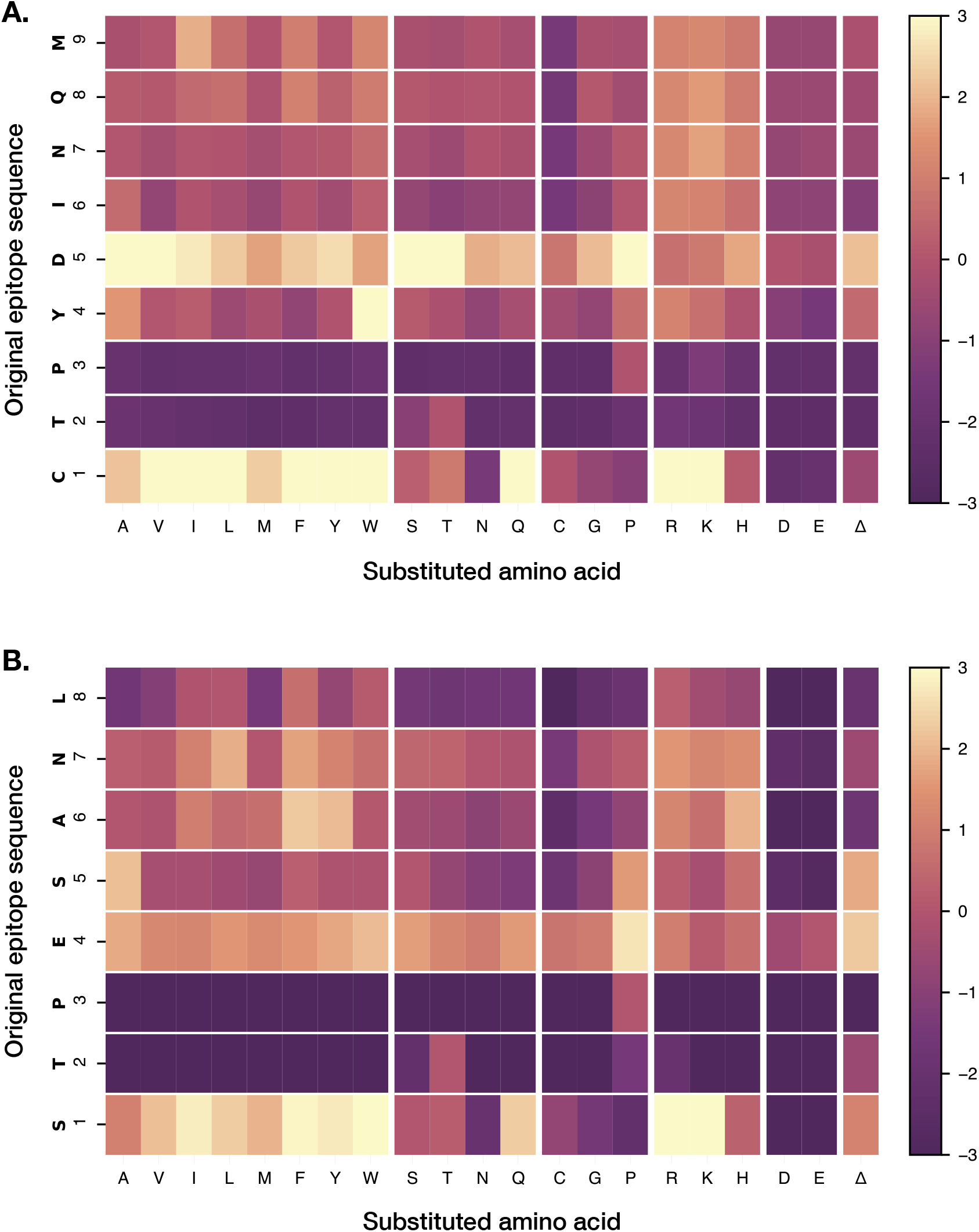
Identification of anchor residues by substitution analysis. Single amino acid substitutions were made at each position of the Gag CM9 (A) and SL8 (B) epitopes. Each box represents a peptide with the indicated substitution, and the fold change in signal intensities for each substitution (compared to the wild-type sequence) is reported. Position numbers are reported along the left axis of the heatmap; the fold change color scale is indicated at right.

### Measuring IFN-γ ELISPOT responses to SIVmac239 peptides that bind to Mamu-A1*001

We assessed whether the MHC peptide array can be used to efficiently screen for putative novel CD8 T cell epitopes by performing IFN-γ ELISPOTs for top-binding peptides. We tested six previously-established Mamu-A1*001 CD8 T cell epitopes plus the top 86 SIVmac239 peptides that bound Mamu-A1*001 with the highest binding scores, using PBMCs from four SIVmac239-infected Mamu-A1*001-positive rhesus macaques. PBMC from each animal ranged from 12 to 47 weeks post infection. Of the established epitopes, Gag CM9 was positive in all four animals; Tat SL8 was positive in two (TVPWPNASL was positive in one, and LGPHYTPKIV, CAPPGYAL, and CAPPGYALL were positive in none). Of the 86 peptides selected from the array, 10 were positive in at least one animal and three were positive in three animals (Supplemental table II). All three of the peptides positive in multiple animals (STPESANLG, STPESANLGE, TPESANLGE) overlapped Tat SL8, and in one animal were positive even though SL8 itself was not. Given the extensive research on Mamu-A1*001-restricted T cell responses to SIVmac239 over the past two decades, it is perhaps not surprising that we did not identify additional shared responses to epitopes in this virus restricted by this allele.

To quantify and confirm these responses, we produced Mamu-A1*001 tetramers folded with Gag CM9, Tat SL8, and the three peptides identified by ELISPOT. We tested these for CD8 T cell binding using PBMCs from two Mamu-A1*001-positive animals whose cells we used in our ELISPOT experiments (one animal that did and one that did not show responses to the three ELISPOT-positive peptides) and one animal that did not have Mamu-A1*001. For Gag CM9, Tat SL8, STPESANLG, and STPESANLGE (but not TPESANLGE), we identified populations of tetramer-positive CD8 T cells within the PBMCs from the Mamu-A1*001-positive, ELISPOT-positive animal, and much smaller populations in those from the Mamu-A1*001-positive, ELISPOT-negative animal (Figure 3). As expected, we did not identify populations of tetramer-positive CD8 T cells in the Mamu-A1*001-negative animal.

**Figure 3:**
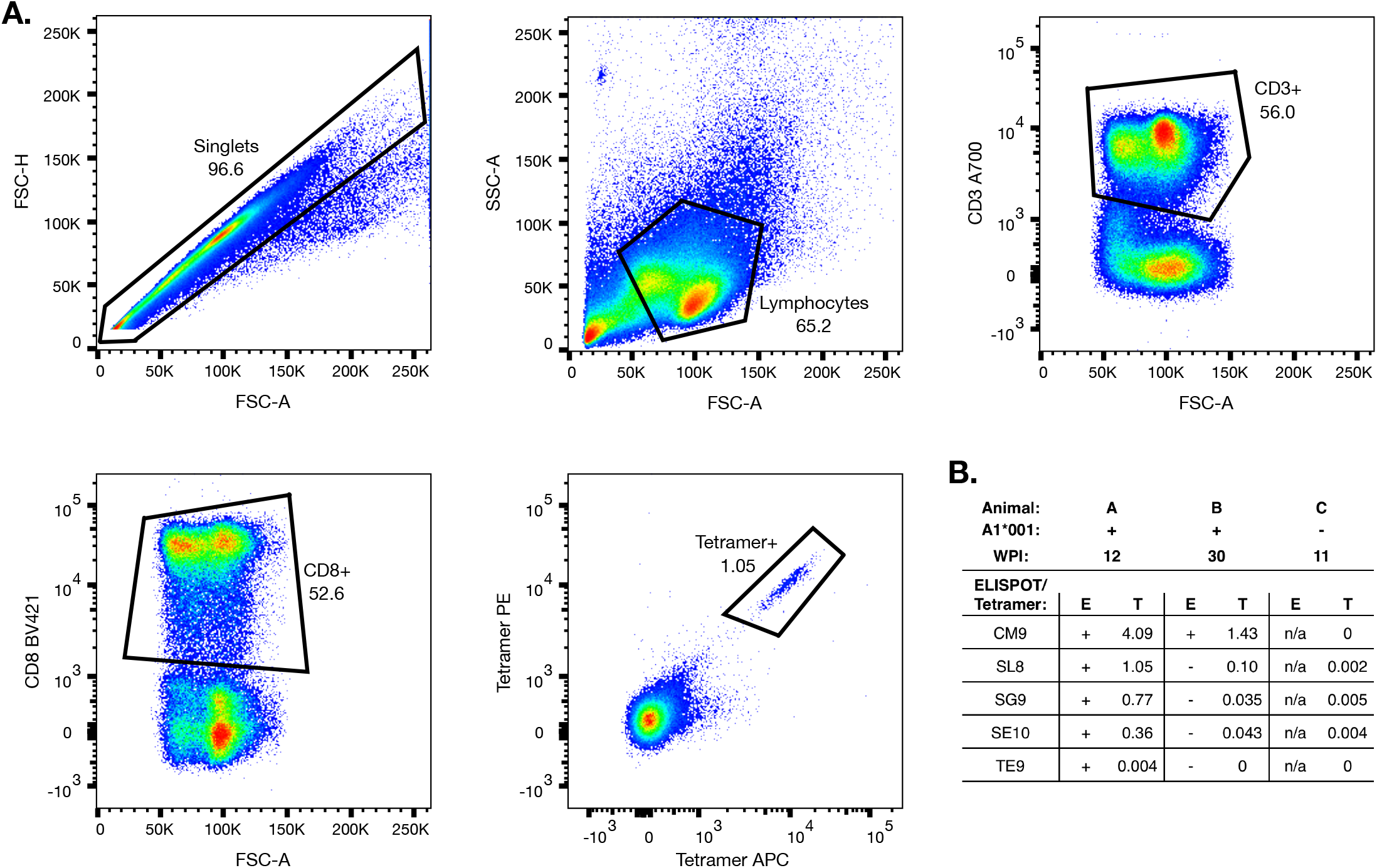
Tetramer analysis. Mamu-A1*001 MHC-peptide tetramers were generated and tested for binding to PBMC from SIV-infected animals. (A) A representative sequential gating strategy for the flow cytometry data is shown. (B) Tetramer binding to PBMC from three animals was tested; shown are the ELISPOT responses to each peptide (STPESANLG, STPESANLGE, and TPESANLG are abbreviated as SG9, SE10, and TE9 respectively), and the percentage of CD8 T cells that were tetramer-positive.

### Comparison of Mamu-A1*001 binding peptides across SIV strains

Due to its large capacity, the peptide array offers the ability to compare binding of MHC molecules to peptides across several related pathogen strains. As a proof of concept, we compared Mamu-A1*001 binding to SIVmac239 and SIVmac251 peptides. Most of the known Mamu-A1*001-restricted SIVmac239 epitopes have the same sequence in SIVmac251, but a few have amino acid variations. Of the six SIVmac239 epitopes that had amino acid variations in SIVmac251, we identified one (QS<u>P</u>GGL<u>D</u>KGL in SlVmac239, QS<u>L</u>GGL<u>G</u>KGL in SIVmac251) that lost its proline in position 3; as we would expect based on the importance of position 3 in binding, this epitope was in the top 192 highest-binding peptides for SIVmac239 but not for SIVmac251 (Supplemental table III).

We additionally compared Mamu-A1*001 binding at the region corresponding to the SIVmac239 Tat SL8 epitope across all viruses on the array. CD8 T cell responses to the SL8 epitope in animals carrying the Mamu-A1*001 allele have been associated with suppression of viral replication, and mutations in this region are commonly implicated in viral escape (26, 27). Identifying whether Mamu-A1*001 is likely to be able to bind to this region despite amino acid sequence differences will be important in determining whether this epitope region is likely to be immunodominant in viral strains other than SIVmac239. SIVmac251 strains, also commonly used in research, are split between having the SL8 sequence and TTPESANL (TL8). TL8 has previously been found to be immunodominant; however, this was first established in experiments studying macaques infected with SIVmac239 – suggesting that there is some latitude in Mamu-A1*001’s ability to bind to that region (13). The peptide array results affirm TL8 as a strong binder to Mamu-A1*001, with a higher binding score (8.72) than SL8 (8.33). We found that all SIVsm (isolated from sooty mangabeys) strains had the sequence PTPESANL (PL8), an escape variant seen in SIVmac239 infection that exhibits reduced binding to Mamu-A1*001 (27). Similarly, on our array we found that PL8 had a lower binding score than SL8 and TL8 (6.28), but one that was still above baseline.

### Correlation of binding scores to IC50 values

IC_50_ values obtained in competitive binding assays are customarily used to represent MHC-peptide binding affinity; however, the equilibrium dissociation constant (k_d_) is more commonly used to quantify protein-ligand binding affinity and is generally preferred as it does not depend on experimental conditions. The peptide array analysis outputs binding scores for each MHC-peptide combination measured in arbitrary fluorescence intensity units, which reflect binding affinity. While we expect these values to correlate with those obtained from competitive binding assays, recent studies have found that IC_50_ values do not accurately estimate kd values, making it challenging to make a direct comparison between IC_50_ and fluorescence intensity units (28). However, using the IC50 data available in the literature for peptides tested for binding to Mamu-A1*001, -A1*002, -B*008, and -B*017, we do find that peptide array binding scores for peptides that were previously found to bind an MHC molecule (IC_50_ ≤ 500 nM) were significantly higher (Median = 0.51; 95% CI = 0.34, 0.70; *p* < 0.001, using a Wilcoxon rank-sum test) than for those that did not bind MHC (IC_50_ > 500 nM) (Figure 4). This dataset had a small selection of binding peptides for all MHC molecules with non-binding peptides reported only for Mamu-B*008; using a dataset of IC_50_ values for all peptide/MHC combinations generated using Stabilized Matrix Method (SMM) MHC-peptide binding prediction software where experimental values were unavailable yielded the same significant difference (data not shown).

**Figure 4:**
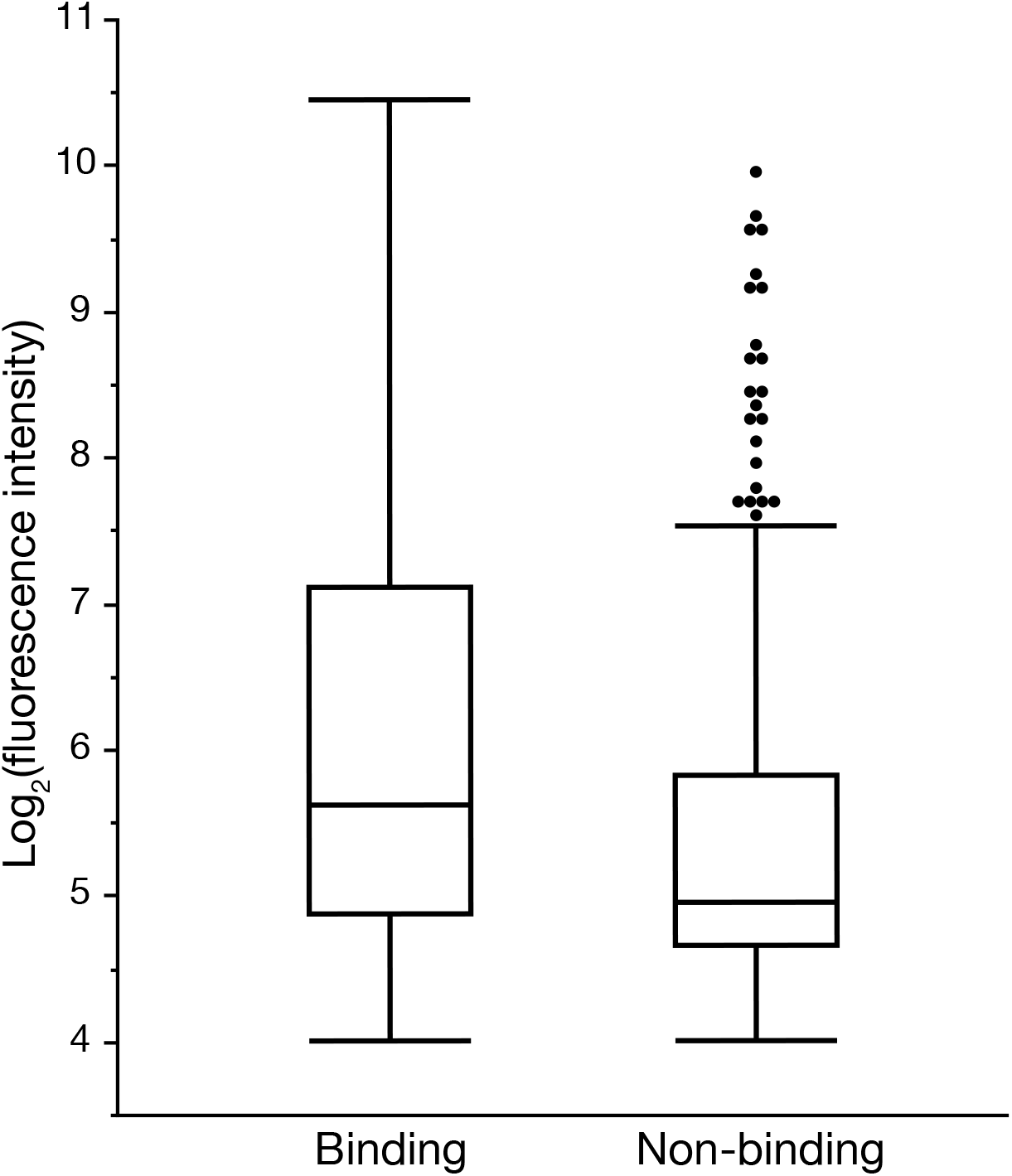
Comparison of IC50 and fluorescence intensity scores. Peptides with experimentally obtained IC50 values for binding to Mamu-A1*001, -A1*002, -B*008, or -B*017 were grouped by IC50 value into binding (IC50 ≤ 500 nM) and non-binding (IC50 > 500 nM) peptides. Binding peptides had significantly higher fluorescence intensity scores than non-binding peptides (*p* < 0.001, using a Wilcoxon rank-sum test).

## Discussion

Identification of CD8 T cell epitopes has historically relied on low-throughput experiments with cumbersome peptide screening methods. In an effort to develop a high-throughput method for screening peptides for MHC binding prior to determining CD8 T cell responses, we used ultradense peptide array technology to test the binding of class I MHC molecules to viral peptides. Compared to conventional methods, ultradense peptide arrays provide massive amounts of information regarding binding of class I MHC molecules to thousands of individual peptides in a single experiment. In this study, of the 86 peptides that bound Mamu-A1*001 with the highest fluorescence intensity scores on the array, we identified 10 peptides that were positive in at least one animal and three that were positive in three animals using IFN-γ ELISPOTs and PBMCs from four SIVmac239-infected Mamu-A1*001-positive rhesus macaques. These peptides were not reported as epitopes previously, thus demonstrating the power of the peptide array method as an alternative approach to screen for putative novel immunogenic epitopes. We additionally demonstrate that many of the same epitopes previously found to bind each molecule and induce CD8 T cell responses were identified as high-binding peptides in our array; 27-64% of the previously-identified epitopes for each MHC molecule were found in the top 192 high-binding peptides on our array.

Despite these initial promising results, many known epitopes were not detected by our method. This may be due in part to the choice of 192 peptides as a convenient cutoff for “high binding.” Because the peptide array binding scores do not directly correspond to IC_50_ values from competitive binding assays, we refrain from choosing a hard binding score cutoff to distinguish binding from non-binding peptides. While there was a significant difference (*p* < 0.001) in fluorescence intensity values between peptides that were previously found to bind or not bind to MHC molecules based on competitive binding assays, there were some instances where fluorescence intensity values did not reflect the results of the competitive binding assays. For example,the Tat SL8 epitope’s high fluorescence intensity (7.15 sd above the median for Mamu-A1*001) and low IC_50_ value both indicate strong peptide binding. However, the Gag CM9 epitope showed a substantially lower fluorescence intensity (2.36 sd above the median for Mamu-A1*001), but also had a low IC_50_ value. These discrepancies may be attributable to differences in how certain peptides or MHC molecules perform on the array; certainly, it would be reasonable to expect that some differences may arise because peptides are anchored on the array rather than free-floating. Unfortunately, neither competitive binding assays nor peptide arrays truly mimic *in vivo* MHC-peptide binding, so without further optimization of the method, it would be premature to label one method as “more true” than the other.

We tested a portion of the high-binding peptides identified here by IFN-γ ELISPOT as a proof of concept. We identified three SIVmac239 peptides that strongly bound Mamu-A1*001 that had positive responses in at least two animals and were not previously identified epitopes; two of the tetramers generated to test these responses were functional. All three peptides overlapped the Tat SL8 epitope, and the two functional tetramers had weaker responses than the SL8 tetramer, which may suggest that these are not separate epitopes but simply extensions of the SL8 minimal optimal epitope. Still, this indicates that a researcher using the array to look for MHC binding to an unknown pathogen’s proteome could reasonably expect to find CD8 T cell epitopes in a subset of high-binding peptides identified by the array; moreover, this demonstrates that, at least in some cases, peptide binding to clusters of related sequences (ie., 8-mer, 9-mer, and 10-mer peptides containing the same component 8-mer) could be used as a filter to identify interesting peptides for follow-up. Using the array as a screening tool could thus drastically reduce the amount of work and resources required to identify novel epitopes.

MHC-peptide array technology has not yet been optimized to work with class II MHC molecules. CD8 T cell responses are not the sole arbiter of protective immunity in HIV vaccination; a better understanding of CD4 responses is necessary for developing a more comprehensive picture of the determinants of a successful vaccine. Assessing CD4 T cell responses in PBMCs and developing class II tetramers is currently more difficult than for CD8 T cells and class I MHC. While this will present challenges in validating a class II array, it underscores the need for a method for rapidly screening for potential CD4 T cell responses, and optimizing a class II array will be an important future direction.

Ultradense peptide arrays offer a method for rapidly mapping class I MHC binding to thousands of individual peptides. While we used peptides derived from SIV and SHIV here, the scalability of ultradense peptide arrays makes them an attractive option for characterizing MHC class I binding to peptides from larger, more complex pathogens. *Mycobacterium tuberculosis* (Mtb) Erdman strain, for instance, has 4,246 annotated protein sequences, in contrast to nine for SIV. The size of the Mtb proteome makes thorough analyses of MHC binding across the full proteome nearly impossible with conventional methods, but just three peptide array chips would be sufficient to accommodate a full 8-, 9-, and 10-mer peptide analysis for a single MHC allele. The ability to screen peptides for MHC binding en masse prior to performing *in vitro* tests will streamline epitope identification studies that might otherwise be prohibitively large and expensive to conduct.

## Acknowledgments

The authors would like to thank Amy Ellis for her help in analyzing the tetramer flow data.

