## Supplemental materials for "High-throughput identification of MHC class I binding peptides using an ultradense peptide array"

**Supplemental table 1:** GenBank accession numbers of SIV and SHIV strains included on the peptide array. All sequences were used without modification; however, the SIVmac239 GenBank entry has a truncated Nef sequence, so a full-length version was manually added.

| Virus | Strain/Isolate | Accession No. | Virus | Strain/Isolate | Accession No. |
|---|---|---|---|---|---|
| SIV | SIVmac239 | M33262.1 | SIV | SIVmacL28-C1 | KU892415.1 |
| SIV | pE660.CG7V | JX648291.1 | SIV | SIVmac746 | KU955513.1 |
| SIV | pE660.CG7G | JX648292.1 | SIV | SIVmac766 | KU955514.1 |
| SIV | SIVmac251.CM.p3c22 | KC522216.1 | SIV | SIVmacCR53 | KU955515.1 |
| SIV | SIVmac251.CM.p3d18 | KC522217.1 | SIV | SIVmacPBE | KU955516.1 |
| SIV | SIVmac251.CM.p3c20 | KC522218.1 | SIV | SIVsmR02012 | KU955517.1 |
| SIV | SIVmac251.CM.tb9 | KC522219.1 | SIV | SIVsmR95117 | KU955518.1 |
| SIV | SIVmac251.CM.p3a6 | KC522220.1 | SIV | SIVsmE660-FL14 | JQ864087.1 |
| SIV | SIVmac251.CM.p3c6 | KC522221.1 | SIV | SIVsmE660-FL8 | JQ864086.1 |
| SIV | SIVmac251.CM.p3d20 | KC522222.1 | SIV | SIVsmE660-FL6 | JQ864085.1 |
| SIV | SIVmac251.CM.p3d1 | KC522223.1 | SIV | SIVsmE660-FL10 | JQ864084.1 |
| SIV | SIVmac251.CM.p3e10 | KC522224.1 | SIV | SIVsmH635FC | DQ201174.1 |
| SIV | SIVmac251.CM.p3c7 | KC522225.1 | SIV | SIVsmH635SB10 | DQ201173.1 |
| SIV | SIVmac251.CM.p3e9 | KC522226.1 | SIV | SIVsmH635F-L3 | DQ201172.1 |
| SIV | SIVmac251.DB.p1e18 | KC522227.1 | SIV | SIVsmE543 | U72748.2 |
| SIV | SIVmac251.DB.p4e16 | KC522228.1 | SIV | SIVmac32H | D01065.1 |
| SIV | SIVmac251.DB.p4g16 | KC522229.1 | SIV | SIVMne027 | U79412.1 |
| SIV | SIVmac251.DB.p1e23 | KC522230.1 | SHIV | SHIV-1157ipd3N4 | DQ779174.2 |
| SIV | SIVmac251.DB.p4a21 | KC522231.1 | SHIV | pSF257.2 | KU521530.1 |
| SIV | SIVmac251.DB.p4e11 | KC522232.1 | SHIV | SHIV-89.6P | U89134.1 |
| SIV | SIVmac251.DB.p1e24 | KC522233.1 | SHIV | SHIV_DH12_CL8 | JN560963.1 |
| SIV | SIVmac251.DB.p4f10 | KC522234.1 | SHIV | SHIV_DH12_CL7 | JN560962.1 |
| SIV | SIVmac251.DB.p1e19 | KC522235.1 | SHIV | SHIV_AD8 | JN560961.1 |
| SIV | SIVmac251.DB.p4b1 | KC522236.1 | SHIV | pSHIV_AD8_B_ES_3N6 | JN560960.1 |
| SIV | SIVmac251.DB.p4e19 | KC522237.1 | SHIV | SHIV AD8 B ES 3T7 | JN560959.1 |
| SIV | SIVmac251.DB.p1e22 | KC522238.1 | SHIV | SHIV-C2/1 | AF217181.1 |
| SIV | SIVmac251.RD.p3h14 | KC522239.1 | SHIV | SHIV-HXBc2P 3.2 | AF041850.1 |
| SIV | SIVmac251.RD.p3i16 | KC522240.1 | SHIV | SHIV-4 | AF038399.1 |
| SIV | SIVmac251.RD.p3g16 | KC522241.1 | SHIV | SHIV-89.6 | AF038398.1 |
| SIV | SIVmac251.RD.p3g21 | KC522242.1 | SHIV | SHIV-89.6Pcy243 | EF672090.1 |
| SIV | SIVmac251.RD.p3j14 | KC522243.1 | SHIV | SHIV.B.YU2C | KU958489.1 |
| SIV | SIVmac251.RD.p3j17 | KC522244.1 | SHIV | SHIV.C.CH848.dCT | KU958488.1 |
| SIV | SIVmac251.RD.p3g14 | KC522245.1 | SHIV | SHIV.C.CH505.dCT | KU958487.1 |
| SIV | SIVmac251.RD.p3j19 | KC522246.1 | SHIV | SHIV.D.191859.dCT | KU958486.1 |
| SIV | SIVmac251.RD.tf20 | KC522247.1 | SHIV | SHIV.D.191859 | KU958485.1 |
| SIV | SIVmac251.RD.p3f16 | KC522248.1 | SHIV | SHIV.A.BG505.dCT | KU958484.1 |
| SIV | SIVmac251.RD.p3i17 | KC522249.1 | SHIV | genomic RNA | AB177846.1 |
| SIV | SIVmac251.RD.p3h1 | KC522250.1 | SHIV | SHIV_SF162 | KF042063.1 |

| **Virus** | **Strain/Isolate** | **Accession No.** |
|---|---|---|
| SIV | SIVmac251.RD.tf1 | KC522251.1 |
| SIV | SIVmac251.RD.p3h4 | KC522252.1 |
| SIV | SIVmac251.RD.tf5 | KC522253.1 |

| **Virus** | **Strain/Isolate** | **Accession No.** |
|---|---|---|
| SHIV | SHIVku2 | AY751799.1 |
| SHIV | 1B3 | AF465242.1 |
| SHIV | NHP model of AIDS | BD161892.1 |

**Supplemental Table II: Summary of Mamu-A1*001 ELISPOT results**

Of the SIVmac239 peptides that strongly bound Mamu-A1*001, we tested the top 86 highest-binding peptides, not including six previously established epitopes (indicated by asterisk), by IFN-γ ELISPOT. Shown here are the peptide array binding scores as well as the previously-established $IC_{50}$ values where available {11014195, 11134287}. We performed IFN-γ ELISPOTs using PBMC from four Mamu-A1*001-positive, SIVmac239-infected rhesus macaques, and the number of animals for which each peptide was positive (bold text) is indicated.

| Peptide array ranking | Sequence | Position | Peptide array binding score (fluorescence intensity units) | $IC_{50}$ (nM) | Number of positive ELISPOTs |
|---|---|---|---|---|---|
| **230** | **CTPYDINQM*** | **Gag 181-189** | **45.79** | **86** | **4** |
| 1621 | LGPHYTPKIV* | Pol 147-156 | 20.42 | 3.0 | 0 |
| 43 | CAPPGYAL* | Env 233-240 | 330.49 | 1.9 | 0 |
| 59 | CAPPGYALL* | Env 233-241 | 216.87 | 5.5 | 0 |
| **41** | **TVPWPNASL*** | **Env 620-628** | **343.76** | **10** | **1** |
| **44** | **STPESANL*** | **Tat 28-35** | **321.58** | **43** | **2** |
| 1 | LTPKWNNETW | Env 628-637 | 1802.18 | - | 0 |
| 2 | ITPIGLAPT | Env 502-510 | 1399.81 | 286 | 0 |
| 3 | SSPPSYFQQT | Env 726-735 | 1382.68 | - | 0 |
| 4 | LTPKWNNE | Env 628-635 | 1294.58 | - | 0 |
| 5 | ITPIGLAPTD | Env 502-511 | 1238.26 | - | 0 |
| 6 | ITPIGLAP | Env 502-509 | 1191.29 | - | 0 |
| **7** | **VTPNYADIL** | **Vif 100-108** | **1133.76** | **-** | **1** |
| 8 | VTPNYADI | Vif 100-107 | 1125.3 | - | 0 |
| **9** | **VTPNYADILL** | **Vif 100-109** | **1087.67** | **-** | **1** |
| 10 | SSPPSYFQQ | Env 726-734 | 1015.23 | - | 0 |
| 11 | LTPKWNNET | Env 628-636 | 971.16 | - | 0 |
| 12 | LTPEKGWLS | Vif 75-83 | 928.64 | - | 0 |
| 13 | SSPPSYFQ | Env 726-733 | 910.88 | - | 0 |
| 14 | LTPEKGWLST | Vif 75-84 | 891.93 | - | 0 |
| 15 | LSPRTLNAWV | Gag 149-158 | 850.86 | - | 0 |
| 16 | MTPAERLI | Pol 961-968 | 815.79 | 77 | 0 |
| 17 | LSPRTLNAW | Gag 149-157 | 799.88 | 355 | 0 |
| 18 | TPEALCDP | Rev 91-98 | 796.59 | - | 0 |
| 19 | MTPAERLINM | Pol 961-970 | 783.83 | - | 0 |
| 20 | LTPEKGWL | Vif 75-82 | 729.27 | - | 0 |
| 21 | MTPAERLIN | Pol 961-969 | 720.31 | - | 0 |
| 22 | FSSPPSYFQQ | Env 725-734 | 646.93 | - | 0 |
| **23** | **TPESANLGE** | **Tat 29-37** | **620.34** | **-** | **3** |
| 24 | FSSPPSYFQ | Env 725-733 | 615.8 | - | 0 |

| Peptide array ranking | Sequence | Position | Peptide array binding score (fluorescence intensity units) | $IC_{50}$ (nM) | Number of positive ELISPOTs |
|---|---|---|---|---|---|
| 25 | FSSPPSYF | Env 725-732 | 599.01 | - | 0 |
| 26 | TPEALCDPTE | Rev 91-100 | 586.52 | - | 0 |
| 27 | SPPSYFQQ | Env 727-734 | 500.54 | - | 0 |
| 28 | STPPLVRL | Pol 625-632 | 498.93 | 26 | 0 |
| 29 | YTPKIVGGIG | Pol 151-160 | 490.87 | - | 0 |
| 30 | LAPVPIPFA | Gag 372-380 | 474.81 | 50 | 0 |
| 31 | YTPKIVGGI | Pol 151-159 | 460.15 | 26 | 0 |
| 32 | LSPRTLNA | Gag 149-156 | 440.78 | - | 0 |
| 33 | MSPSYVKY | Vpx 62-69 | 435.4 | - | 0 |
| 34 | TPESANLG | Tat 29-36 | 423.46 | - | 0 |
| 35 | STPPLVRLV | Pol 625-633 | 408.71 | 8.7 | 0 |
| 36 | APVPIPFA | Gag 373-380 | 396.06 | - | 0 |
| **37** | **STPESANLGE** | **Tat 28-37** | **352.69** | **-** | **3** |
| 38 | STPPLVRLVF | Pol 625-634 | 347.14 | - | 0 |
| 39 | MSPSYVKYR | Vpx 62-70 | 346.43 | - | 0 |
| 40 | TPESANLGEE | Tat 29-38 | 344.9 | - | 0 |
| 42 | TPAERLIN | Pol 962-969 | 331.75 | - | 0 |
| 45 | TPEALCDPT | Rev 91-99 | 314.9 | - | 0 |
| 46 | LAPVPIPFAA | Gag 372-381 | 314.16 | - | 0 |
| 47 | LAPVPIPF | Gag 372-379 | 304.74 | 29 | 0 |
| 48 | APVPIPFAA | Gag 373-381 | 303.04 | - | 0 |
| **49** | **YSFPDPPTDT** | **Rev 58-67** | **272.97** | **-** | **1** |
| 50 | SPPSYFQQT | Env 727-735 | 269.42 | - | 0 |
| 51 | MSPSYVKYRY | Vpx 62-71 | 268 | - | 0 |
| 52 | VPWPNASL | Env 621-628 | 260.92 | - | 0 |
| 53 | TTVPWPNASL | Env 619-628 | 260.71 | - | 0 |
| 54 | VPWPNASLTP | Env 621-630 | 260.28 | - | 0 |
| **55** | **STPESANLG** | **Tat 28-36** | **247.33** | **-** | **3** |
| 56 | YSFPDPPT | Rev 58-65 | 232.86 | - | 0 |
| 57 | LSPLCITM | Env 104-111 | 228.85 | - | 0 |
| 58 | TPLDLAIQ | Rev 67-74 | 225.71 | - | 0 |
| 60 | VPIPFAAA | Gag 375-382 | 224.98 | - | 0 |
| 61 | TVPWPNAS | Env 620-627 | 205.87 | - | 0 |
| 62 | TPAERLINM | Pol 962-970 | 203.9 | - | 0 |
| 63 | VPWPNASLT | Env 621-629 | 203.81 | - | 0 |
| **64** | **ISTPPLVRL** | **Pol 624-632** | **203.39** | **-** | **1** |

| Peptide array ranking | Sequence | Position | Peptide array binding score (fluorescence intensity units) | $IC_{50}$ (nM) | Number of positive ELISPOTs |
|---|---|---|---|---|---|
| **65** | **QTNPYPTGPG** | **Rev 22-31** | **197.5** | **-** | **1** |
| **66** | **ISTPPLVRLV** | **Pol 624-633** | **192.71** | **-** | **1** |
| 67 | SPRTLNAW | Gag 150-157 | 189.3 | - | 0 |
| 68 | QTNPYPTGP | Rev 22-30 | 188.78 | - | 0 |
| 69 | TPIGLAPT | Env 503-510 | 186.3 | - | 0 |
| 70 | TVPWPNASLT | Env 620-629 | 184.3 | - | 0 |
| 71 | SPAIFQYT | Pol 364-371 | 181.73 | - | 0 |
| 72 | SPPSYFQQTH | Env 727-736 | 179.75 | - | 0 |
| 73 | TPAERLINMI | Pol 962-971 | 178.34 | - | 0 |
| 74 | VPIPFAAAQ | Gag 375-383 | 163.22 | - | 0 |
| 75 | LTACQGVGGP | Gag 348-357 | 162.84 | - | 0 |
| 76 | TPLDLAIQQ | Rev 67-75 | 161.61 | - | 0 |
| 77 | YSFPDPPTD | Rev 58-66 | 160.34 | - | 0 |
| 78 | VFSSPPSYFQ | Env 724-733 | 153.61 | - | 0 |
| 79 | SPAIFQYTM | Pol 364-372 | 152.9 | - | 0 |
| 80 | TPINIFGRN | Pol 189-197 | 152.2 | - | 0 |
| 81 | QTNPYPTG | Rev 22-29 | 152.09 | - | 0 |
| 82 | TPIGLAPTD | Env 503-511 | 151.3 | - | 0 |
| 83 | VPIPFAAAQQ | Gag 375-384 | 150.08 | - | 0 |
| 84 | TTVPWPNAS | Env 619-627 | 150.01 | - | 0 |
| 85 | VFSSPPSYF | Env 724-732 | 148.15 | - | 0 |
| 86 | APPGYALL | Env 234-241 | 147.58 | - | 0 |
| 87 | SLTPKWNNET | Env 627-636 | 145.53 | - | 0 |
| 88 | IPPSRSML | Vpr 94-101 | 145.21 | - | 0 |
| **89** | **NTPEALCDPT** | **Rev 90-99** | **144.86** | **-** | **1** |
| 90 | LSPLCITMR | Env 104-112 | 143.34 | - | 0 |

**Supplemental table III:** Comparison of Mamu-A1*001 binding to known SIVmac239 epitopes and the equivalent SIVmac251 sequences. Sequences with amino acid changes between viruses are indicated in bold. Only one sequence was considered “high binding” in SIVmac239 but not in SIVmac251 as the result of a position three proline to leucine substitution.

| SIVmac239 epitope | SIVmac239 intensity | SIVmac239 top 192 | SIVmac251 sequence | SIVmac251 intensity | SIVmac251 top 192 |
|---|---|---|---|---|---|
| ITPIGLAPT | 10.45 | Yes | ITPIGLAPT | 10.45 | Yes |
| SSPPSYFQQT | 10.43 | Yes | SSPPSYFQQT | 10.43 | Yes |
| **VTPNYADIL** | **10.15** | **Yes** | **VTPDYADIL** | **10.04** | **Yes** |
| **VTPNYADI** | **10.14** | **Yes** | **VTPDYADI** | **9.52** | **Yes** |
| **VTPNYADILL** | **10.09** | **Yes** | **VTPDYADILL** | **9.80** | **Yes** |
| MTPAERLI | 9.67 | Yes | MTPAERLI | 9.67 | Yes |
| LSPRTLNAW | 9.64 | Yes | LSPRTLNAW | 9.64 | Yes |
| **LTPEKGWL** | **9.51** | **Yes** | **LTPERGWL** | **9.38** | **Yes** |
| STPPLVRL | 8.96 | Yes | STPPLVRL | 8.96 | Yes |
| YTPKIVGGI | 8.85 | Yes | YTPKIVGGI | 8.85 | Yes |
| STPPLVRLV | 8.67 | Yes | STPPLVRLV | 8.67 | Yes |
| TVPWPNASL | 8.43 | Yes | TVPWPNASL | 8.43 | Yes |
| CAPPGYAL | 8.37 | Yes | CAPPGYAL | 8.37 | Yes |
| **STPESANL** | **8.33** | **Yes** | **TTPESANL** | **8.72** | **Yes** |
| LAPVPIPF | 8.25 | Yes | LAPVPIPF | 8.25 | Yes |
| CAPPGYALL | 7.76 | Yes | CAPPGYALL | 7.76 | Yes |
| **ASTPESANL** | **6.71** | **Yes** | **ATTPESANL** | **7.43** | **Yes** |
| **QSPGGLDKGL** | **6.46** | **Yes** | **QSLGGLGKGL** | **4.26** | **No** |
| VVPGFQAL | 6.21 | Yes | VVPGFQAL | 6.21 | Yes |
| GSPAIFQYTM | 6.21 | Yes | GSPAIFQYTM | 6.21 | Yes |
| QVPSLQYLA | 6.04 | Yes | QVPSLQYLA | 6.04 | Yes |
| GSPAIFQYT | 5.91 | Yes | GSPAIFQYT | 5.91 | Yes |
| CTPYDINQM | 5.52 | No | CTPYDINQM | 5.52 | No |
| YVPCHIRQI | 4.85 | No | YVPCHIRQI | 4.85 | No |
| GPPPPPPPGL | 4.83 | No | GPPPPPPPGL | 4.83 | No |
| QVPKFHLPV | 4.77 | No | QVPKFHLPV | 4.77 | No |
| IYPGIKTKHL | 4.53 | No | IYPGIKTKHL | 4.53 | No |
| RIPERLERW | 4.50 | No | RIPERLERW | 4.50 | No |
| QNPIPVGNI | 4.41 | No | QNPIPVGNI | 4.41 | No |

| | | |
|---|---|---|
| LGPHYTPKIV | 4.35 | No |
| DPPTNTPEAL | 4.22 | No |
| HLPRELIFQV | 4.20 | No |
| VNPTLEEMLT | 4.05 | No |


| | | |
|---|---|---|
| LGPHYTPKIV | 4.35 | No |
| DPPTNTPEAL | 4.22 | No |
| HLPRELIFQV | 4.20 | No |
| VNPTLEEMLT | 4.05 | No |
